# Compositional clustering in task structure learning

**DOI:** 10.1101/196923

**Authors:** Nicholas Franklin, Michael J. Frank

## Abstract

Humans are remarkably adept at generalizing knowledge between experiences in a way that can be difficult for computers. Often, this entails generalizing constituent pieces of experiences that do not fully overlap, but nonetheless share useful similarities with, previously acquired knowledge. However, it is often unclear how knowledge gained in one context should generalize to another. Previous computational models and data suggest that rather than learning about each individual context, humans build latent abstract structures and learn to link these structures to arbitrary contexts, facilitating generalization. In these models, task structures that are more popular across contexts are more likely to be revisited in new contexts. However, these models can only re-use policies as a whole and are unable to transfer knowledge about the transition structure of the environment even if only the goal has changed (or vice-versa). This contrasts with ecological settings, where some aspects of task structure, such as the transition function, will be shared between context separately from other aspects, such as the reward function. Here, we develop a novel non-parametric Bayesian agent that forms independent latent clusters for transition and reward functions, affording separable transfer of their constituent parts across contexts. We show that the relative performance of this agent compared to an agent that jointly clusters reward and transition functions depends environmental task statistics: the mutual information between transition and reward functions and the stochasticity of the observations. We formalize our analysis through an information theoretic account of the priors, and propose a meta learning agent that dynamically arbitrates between strategies across task domains to optimize a statistical tradeoff.

**Author summary:** A musician may learn to generalize behaviors across instruments for different purposes, for example, reusing hand motions used when playing classical on the flute to play jazz on the saxophone. Conversely, she may learn to play a single song across many instruments that require completely distinct physical motions, but nonetheless transfer knowledge between them. This degree of compositionality is often absent from computational frameworks of learning, forcing agents either to generalize entire learned policies or to learn new policies from scratch. Here, we propose a solution to this problem that allows an agent to generalize components of a policy independently and compare it to an agent that generalizes components as a whole. We show that the degree to which one form of generalization is favored over the other is dependent on the features of task domain, with independent generalization of task components favored in environments with weak relationships between components or high degrees of noise and joint generalization of task components favored when there is a clear, discoverable relationship between task components. Furthermore, we show that the overall meta structure of the environment can be learned and leveraged by an agent that dynamically arbitrates between these forms of structure learning.

## Introduction

Compared to artificial agents, humans exhibit remarkable flexibility in our ability to rapidly, spontaneously and appropriately learn to behave in unfamiliar situations, by generalizing past experience and performing symbolic-like operations on constituent components of knowledge [1]. Formal models of human learning have cast generalization as an inference problem in which people learn a shared (latent) task structure across multiple contexts and then infer which causal structure best suits the current scenario [2,3]. In these models, a context, typically an observable (or partially observable) feature of the environment, is linked to a learnable set of task statistics or rules. Based on statistics and the opportunity for generalization, the learner has to infer which environmental features (stimulus dimensions, episodes, etc.) should constitute the context that signals the overall task structure, and, simultaneously, which features are indicative of the specific appropriate behaviors for the inferred task structure. This learning strategy is well captured by Bayesian nonparametric models, and neural network approximations thereof, that impose a hierarchical clustering process onto learning task structures [3,4]. A learner infers the probability that two contexts are members of the same task cluster via Bayesian inference, and in novel situations, has a prior to reapply the task structures that have been more popular across disparate contexts, while also allowing for the potential to create a new structure as needed. Empirical studies have provided evidence that humans spontaneously impute such hierarchical structure, which facilitates future transfer, whether or not it is immediately beneficial – and, indeed, even if it is costly – to initial learning [3–5].

These clustering models can account for aspects of human generalization that are not well explained by standard models of learning. This approach to generalization, treating multiple contexts as sharing a common task structure, is similar to artificial agents that reuse previously learned policies in novel tasks when the statistics are sufficiently similar [6–9]. However, a key limitation to these clustering models of generalization is that policies of the agent are generalized as a unit. That is, in a new context, a previously learned policy can either be reused or a new policy must be learned from scratch. This can be problematic as policies are often not robust to untrained variation in task structure [10–12]. Thus, a previously learned policy can lead to a poor outcome in a new context even if there is a substantial degree of shared structure.

Because task structures are either reused or not as a whole, the ability to reuse and share component parts of knowledge is limited; that is, they are not *compositional*. Compositionality, or the ability to bind (compose) information together in a rule governed way, has long been thought to be a core aspect of human cognition [1,13]. Importantly, ecological contexts often share a partial structure, limiting the applicability of previously learned policies but nonetheless providing a generalization advantage to a compositional agent.

To provide a naturalistic example, an adept musician can transfer a learned song between a piano and a guitar, even as the two instruments require completely different physical movements, implying that goals can be generalized and reused independently of the actions needed to achieve them. A clustering model that generalizes entire task structures cannot account for this behavior, and would require instead that an agent would need to relearn a song from scratch to play it on a new instrument. Worse, this clustering scheme would predict an unlikely interference effect where the similar outcome of playing the same song on two instruments results in the model incorrectly pooling motor policies across instruments.

Here, we propose a framework to address one aspect of compositionality by decomposing task structures – and their separable potential for clustering – into reward functions and transition functions. These two independent functions of a Markov decision process are suitable units of generalization: if we assume that an agent has knowledge of a state-space and the set of available actions, then the reward and transition functions are sufficient to determine the optimal policy. In real-world scenarios, a reward function may correspond to the objective of an agent (what it would like to achieve and the environmental states that produce these goals). A transition function determines how the agent’s actions affect its environment (i.e., the subsequent states). For example, when playing music a reward function might correspond to the desired sequence of notes (a scale, or a song) while the transition function might correspond to the actions needed to produce notes on an instrument. When picking up a new form of guitar, it may be sufficient for a musician to play one or two strings which may then afford inference of the entire transition functions (the tuning: strings and frets needed to obtain each note). Here, we are concerned with how the inference of one (reward or transition) function affects generalization of the other.

We consider two approaches to clustering and compare their relative generalization advantages as a function of environmental statistics. The *independent clustering* agent supports generalization by clustering contexts into independent sets defined by the reward and transition statistics, respectively. In contrast, the *joint clustering* agent clusters contexts into a single set of clusters that binds together the transition and reward functions (hence amounting to previous models of task-set structure that cluster and re-use policies [3-5]). Necessarily, independent clustering is compositional and requires the binding of two independent functions.

We show that these two models lead to different predictions depending on the task environment, and we provide an information theoretic analysis to formalize and quantify the bounds of these advantages/disadvantages. In environments where there is a clear, discoverable relationship between transitions and rewards, joint clustering facilitates generalization by allowing an agent to infer one function based on observations that are informative about the other. Nonetheless, we show that independent clustering can lead to superior generalization even in such cases when the transition-reward relationship is weak, difficult to discover, or costly to do so. Finally, we develop a meta-structure learning agent that can infer whether the overall environment is better described by independent or joint statistics.

## Models

To provide a test-bed for characterizing the effects compositional structure, we consider a series of navigation tasks by utilizing grid worlds as a simplification of real-world environments. In these grid worlds, an agent learns to navigate by learning transition functions (the consequences of its actions in terms of subsequent states) and separately learns a reward function (the reward values of locations, or goals) as it navigates. At each point in time, the agent is given a state tuple *s* =< *x*,*c* > where 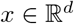 is a vector of state variables (for example, a location vector in coordinate space) and 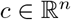 is a context vector. Here, we define “context” as a vector denoting some mutable property of the world (for example, the presence or absence of rain, an episodic period of time, etc.) that constrain the statistics of the task domain, whereby these task statistics are consistent for each context that cues the relevant task. Formally, for each context we can define a Markov decision process (MDP) with state variables *x* ∈ *X*, actions *a* ∈ *A*, a reward function mapping state variables and actions to a real valued number 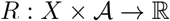, and a transition function mapping state variables and actions to a probability distribution over successors 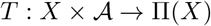 II(X).

For the purpose of simplicity, we assume that the agent knows the spatial relationship between states (i.e., it has access to a spatial map of its current position and adjacent positions) but has to learn how its actions take it from one state to another. Specifically, we assume the agent knows a set of cardinal movements 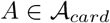, where each cardinal movement is a vector that defines a change in the state variables with regard to the known spatial structure per unit time. (For example, in a two dimensional grid world we can define North = <*dx*/*dt*, *dy*/*dt*> = <0,1> as a cardinal movement). We can thus define a transition function in terms of cardinal movements *T*(*x*, *A*, *x*′|*c*) = Pr(*x*′|*x*, *A*, *c*) and cast the navigation problem as the learning of some function that maps primitive actions to cardinal movements 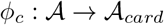, which we assume to be independent of location.

This simplifying assumption has the benefit of providing a model of human navigation, whom we assume understand spatial structure. Note that the function mapping motor actions onto cardinal movements can depend on environmental conditions, and thus, context (for example, wind condition can change the relationship between primitive actions and movements in space for an aerial drone). A similar mapping between arbitrary button presses and movements in the “finger sailing task has been used to provided evidence for model-based action planning in human subjects [14,15]. Similarly, we can express the reward function in terms of cardinal movements based on a location in space, *R_c_*(*x*, *A*) = Pr(*r*|*x, A, c*). This allows us to consider how the agent receives reward as it moves through coordinate space (as opposed to how it receives reward as a function of its actions). Alternatively, we can express the reward function as *R*(*x, x′*) or more simply as *R*(*x*′). The key assumption here is that the reward function is not a function of the agent’s actions but is a function of the consequences of those actions.

The task of the agent is to generate a policy (a function mapping state variables to primitive actions, for each context; 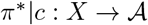 that maximizes its expected future discounted reward [16]. Given a known transition function and reward function, the optimal policy given this task can be defined as:

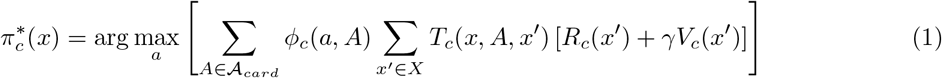

where *V_c_*(*x*) is the optimal value function is defined by the Bellman equation:

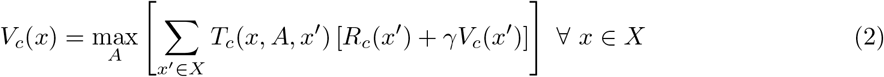

As the relationship between locations in space, *T_c_*, is known to the agent, it is sufficient to learn the cardinal *mapping function ∅_c_*(*a, A*) and *reward function R_c_*(*x, A*) to determine an optimal policy.

While the optimal policy is dependent on both the mapping function and reward function, crucially, the optimal value function is not: it is dependent only on the reward function (and the known transition function *T_c_*). Consequently, an agent can determine an optimal policy as a function of movements through space:

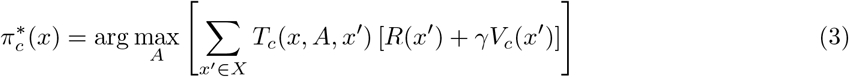

This allows the agent to learn how it can take an action to move through space – the mapping function *∅_c_*(*a, A*) – independently from the desirability of the consequences of these moves *R_c_*(*x*′). This distinction allows for compositionality during generalization, as we will discuss in the following section.

### Context clustering as generalization

A common strategy to support task generalization is to cluster contexts together, assuming they share the same task statistics, if doing so leads to an acceptable degree of error [7]. This logic underlies models of animal Pavlovian learning and transfer [2], human instrumental learning and transfer [3, 4], and category learning [17,18]. Clustering models of human generalization typically rely on a non-parametric Dirichlet process, commonly known as the Chinese restaurant process (CRP), which acts as a clustering prior in a Bayesian inference process. Used in this way, the CRP enforces popularity-based clustering to partition observations, so that the agent will be most likely to reuse those tasks that have been most popular across disparate contexts (as opposed to across experiences; [4]), and has the attractive property of being a non-parametric model that grows with the data [19]. Consequently, it is not necessary to know the number of partitions a priori and the CRP will tend to parsimoniously favor a smaller number of partitions.

As in prior work, we model generalization as the process of inferring the assignment of contexts *k* = {*c*_1:*n*_} into clusters that share common task statistics. But here, we decompose these task statistics to consider the possibility that that all contexts *c* ∈ *k* share either the same reward function and/or mapping function, such that *R_k_*(*x, A*) = *R_c_*(*x, A*) ∀ *c* ∈ *k* and/or *∅_k_*(*a, A*) = *∅_c_*(*a, A*) ∀ *c* ∈ *k*. (We return to the “and/or distinction, which affects whether clustering is independent or joint across reward and mapping functions, in the following section). Formally, we define generalization as the inference

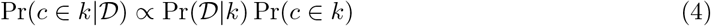

where Pr(*D*|*k*) is the likelihood of the observed data *D* given cluster k, and Pr(*c* ∈ *k*) is a prior over the clustering assignment. As in previous models of generalization, we use the CRP as the cluster prior. If contexts {*c*_1:*n*_} are clustered into *N* ≤ *n* clusters, then the prior probability for any new context *c*_*n*+1_ ∉ {*c*_1:*n*_} is:

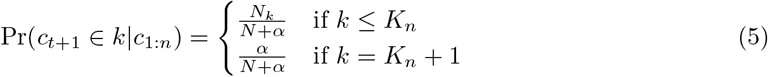

where *N_k_* is the number of contexts associated with cluster *k* and *K_n_* is the number of unique clusters associated with the *n* observed contexts. If *k* ≤ *K_n_*, then *k* is a previously encountered cluster, whereas if *k* = *K_n_* + 1, then *k* is a new cluster. The parameter *α* governs the propensity to assign a new context to a new cluster, that is to create a new task. Higher values of *α* lead to a greater prior probability that a new cluster is created and favors a more expanded task space overall, leading to reduced likelihood of reusing old tasks. Thus, the prior probability that a new context is assigned to an old cluster is proportional to the number of contexts in that cluster (popularity), and the probability that it is assigned to a new cluster is proportional to *α*. As a non-parametric generative process, the prior allows the number of clusters to grow as new contexts are observed. This process is exchangeable, and as such, the order of observation does not alter the inference of the agent [19], though approximate inference algorithms can induce order effects.

### Independent and joint clustering

As we noted above, there are two key functions the agent learns when navigating in a context: *∅_c_*(*a, A*) and *R_c_*(*x, A*). These functions imply that the agent could cluster *∅_c_*(*a, A*) and *R_c_*(*x, A*) jointly, or it could cluster them independently, such that it learns the popularity of each marginalizing over the other. Formally, the *independent clustering* agent [Fig 1, left] assigns each context *c* into two clusters via Bayesian inference as in [4] and using the CRP prior for each cluster [5]. The likelihood function for the two assignments are the mapping and reward functions

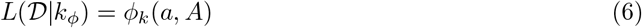

and

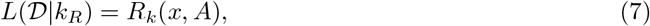

respectively. Conversely, the *joint clustering* agent [Fig 1, right] assigns each context *c* into a single cluster *k* via Bayesian inference [4] and, like the independent clustering agent, uses the CRP as the prior over assignments [5]. The likelihood function for the context-cluster assignment is the product of the mapping and reward functions

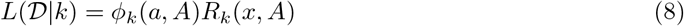

**Fig 1.**
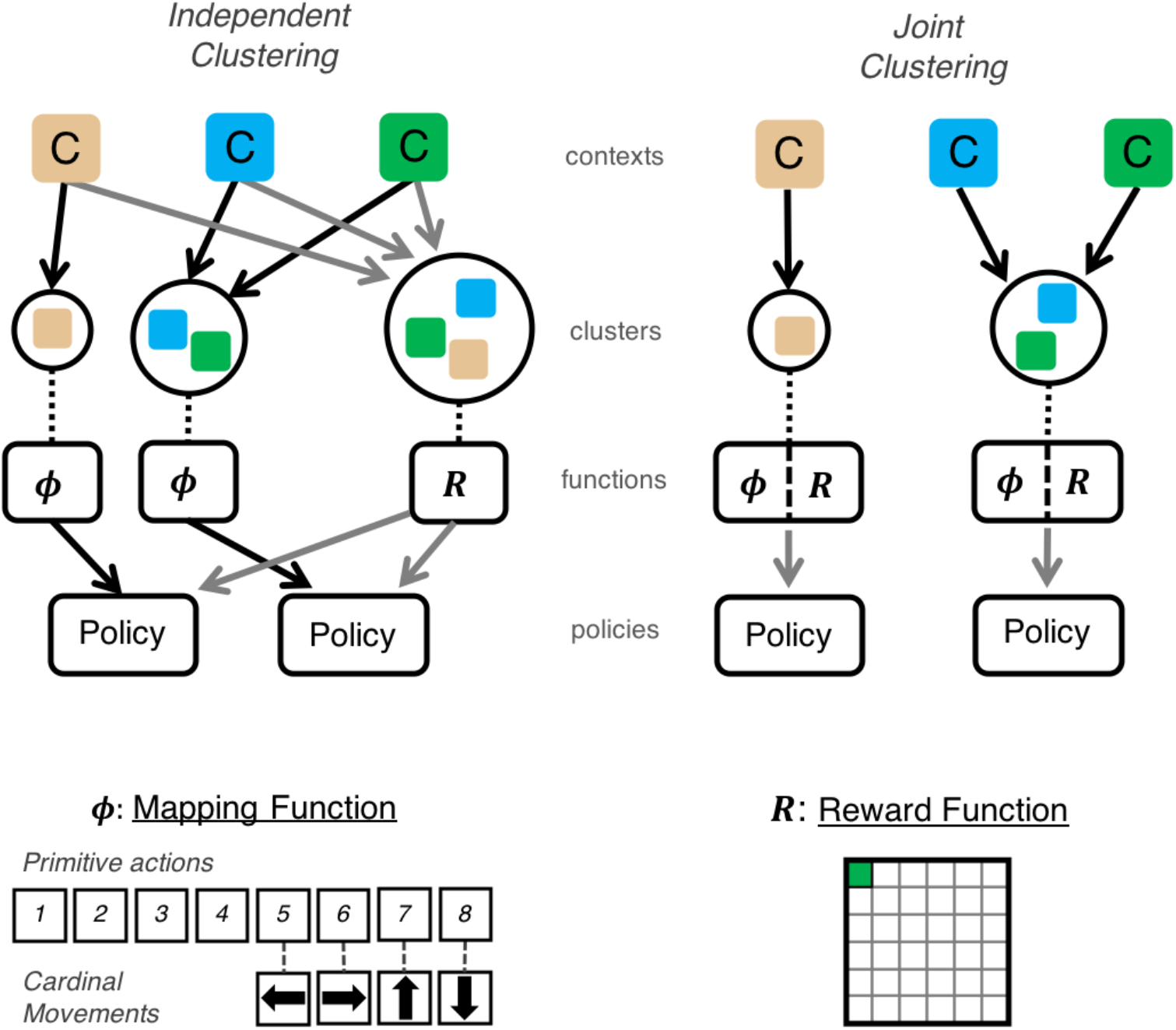
Schematic example of independent and joint clustering agents. *Top Left*: The independent clustering agent groups each context into two clusters, associated with a reward (*R*) and mapping (*∅*) function, respectively. Planning involves combining these functions to generate a policy. The clustering prior induces a parsimony bias such that new contexts are more likely to be assigned to more popular clusters. Arrows denote assignment of context into clusters and creation of policies from component functions. *Top Right*: The joint clustering agent assigns each context into a cluster linked to both functions (i.e., assumes a holistic task structure), and hence the policy is determined by this cluster assignment. In this example, both agents generate the same two policies for the three contexts but the independent clustering agent generalizes the reward function across all three contexts. Bottom: An example mapping (*left*) and reward function (*right*) for a gridworld task.

The joint clustering agent is highly similar to previous non-compositional models of task structure learning and generalization, which have previously been shown to account for human behavior (but without specifically assessing the compositionality issue) [3,4]. For the purpose of comparison with the independent clustering agent, here the joint clustering agent separately learns the functions *R* and *∅*. Previous models, in contrast, have learned model-free policies directly. In the general case, joint clustering does not require the separate representation of policy components (*R* and *∅*) nor does it require a binding operation in the form of planning. However, independent clustering does require the separate representation of policy components that must be bound via planning. Hence, one potential difference between the two approaches is algorithmic complexity, where joint clustering may permit a less complex and computationally costly learning algorithm than independent clustering. In the simulations below, we have equated the agents for algorithmic complexity and examine how inferring the reward and transition functions separately or together affect performance across task domains.

## Results

We first consider two minimal sets of simulations to illustrate the complementary advantages afforded by the two sorts of clustering agents depending on the statistics of the task domain, using a common set of parameters. A third simulation explores how the benefits of compositional clustering can compound in a modified form of the *Rooms* problems previously used to motivate hierarchical RL approaches [20,21]. We show that our independent clustering model with hierarchical decomposition of the state space can facilitate more rapid transfer than that afforded by standard approaches. Finally we conduct an information theoretic analysis to formalize the more general conditions under which each scheme is more beneficial. Code for all of the simulations presented here have been made available in our GitHub repository: https://github.com/nicktfranklin/IndependentClusters

### Simulation 1: Independence of Task Statistics

In the first set of simulations, we simulated a task domain in which four contexts involving different combinations of reward and transition functions. In every trial, a “goal location was hidden in a 6×6 grid world. Agents were randomly placed in the grid world and explored action selection until the goal was reached, at which point the trial ended and the next trial began, with the agent again randomized to a new location. The agent’s task is to find the goal location (encoded as +1 reward for finding the goal and a 0 reward otherwise) as quickly as possible. The agents had a set of eight actions available to them *A* = {*a*_1_, …*a*_8_}, which could be mapped onto one of four cardinal movements *A_card_* = {North, South, East, West}. The agents were exposed to four trials, in which goal locations and mappings were stationary, for each context. Each of the four contexts had a unique combination of one of two goal locations and one of two mappings [Fig 2, A], and hence knowledge about the reward function or mapping function for any context was not informative about the other function. However, because the reward and mapping functions were each common across two contexts, the independent clustering agent can leverage the structure to improve generalization without being bound to the joint distribution of mappings and rewards.

**Fig 2.**
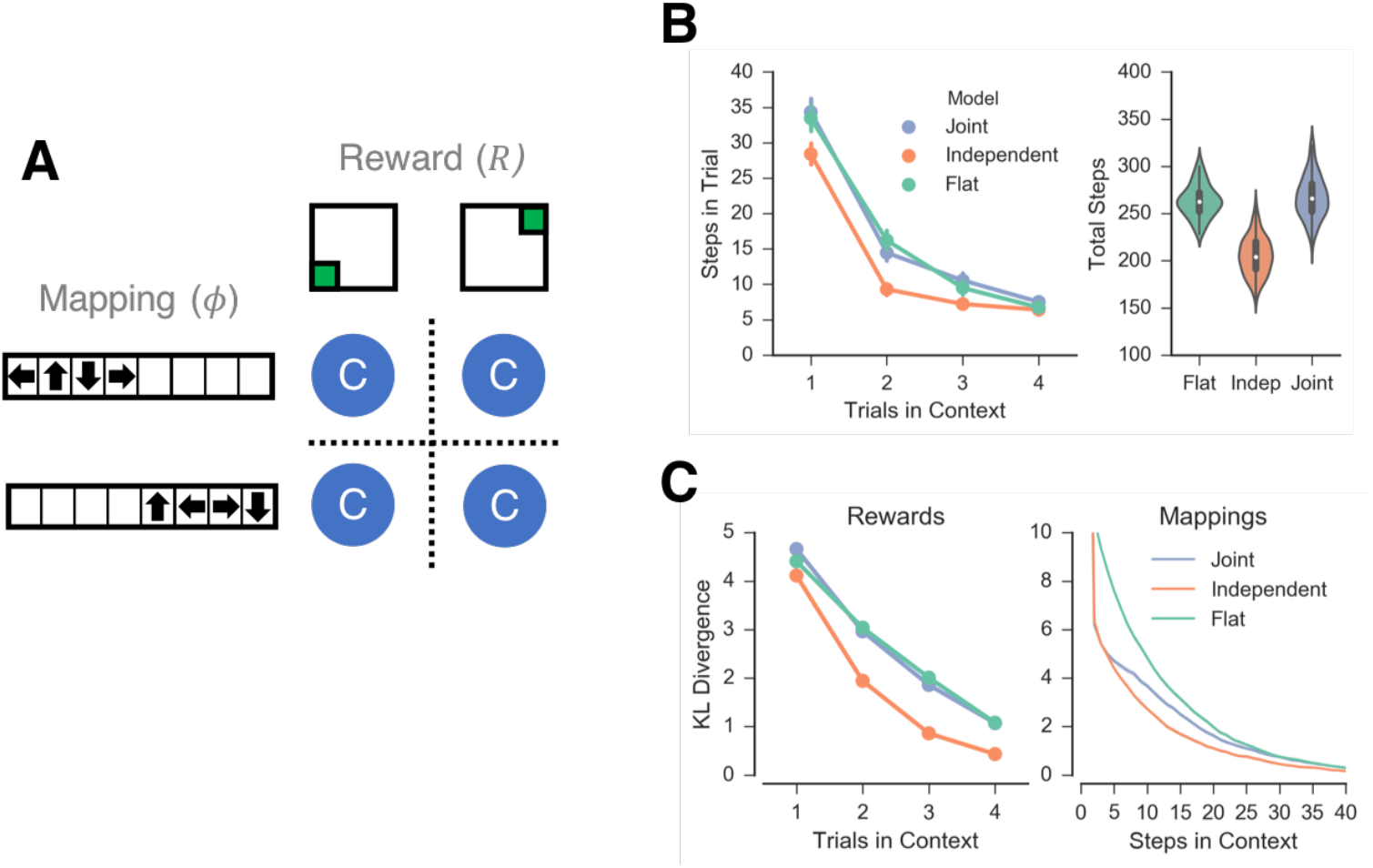
Simulation 1. *A*: Schematic representation of the task domain. Four contexts (blue circles) were simulated, each paired with a unique combination of one of two goal locations (reward functions) and one of two mappings. *B*: Number of steps taken by each agent shown across trials within a single context (left) and over all trials (right). Fewer steps reflect better performance. *C*: KL-divergence of the models’ estimates of the reward (left) and mapping (right) functions as a function of time. Lower KL-divergence represents better function estimates. Time shown as the number of trials in a context (left) and the number of steps in a context collapsed across trials (right) for clarity.

In addition to the independent and joint clustering agents, for comparison, we also simulated a “flat (non-hierarchical decomposition of “context and “state) agent that does not cluster contexts at all and hence has to learn anew in each context. (The flat agent is a special case of both the independent and joint clustering agents such that *k_i_* = {*c_i_*} ∀ *i*). We used hypothesis-based inference, where each hypothesis comprised a proposal assignment of contexts in to clusters, *h*: *c* ∈ *k*, defined generatively, such that when a new context is encountered the hypothesis space is augmented. For each hypothesis, maximum likelihood estimation was used to generate the estimates 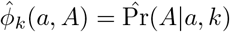 and 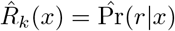. To encourage optimistic exploration, 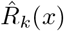 was initialized to the maximum observable reward (Pr(*r*|*x*) = 1) with a low confidence prior using a conjugate beta distribution of Beta(0.01, 0).

The belief distribution over the hypothesis space is defined by the posterior probability of the clustering assignments [4]. Calculating the posterior distribution over the full hypothesis space is computationally intractable, as the size of the hypothesis space grows combinatorially with the number of contexts. As an approximation, we pruned hypotheses with small probability (less than 1/10x posterior probability of the maximum *a posteriori* (MAP) hypothesis) from the hypothesis space. We further approximated the inference problem by using the MAP hypothesis, rather than sampling from the entire distribution, during action selection [3,22]. Value iteration was used to solve the system of equations defined by [3] using the values of 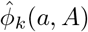 and 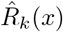 associated with the MAP hypothesis(es). A state-action value function, defined here in terms of cardinal movements was used with a softmax action selection policy to select cardinal movements:

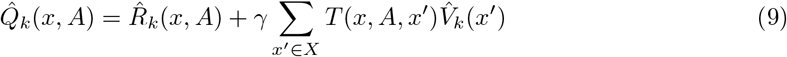

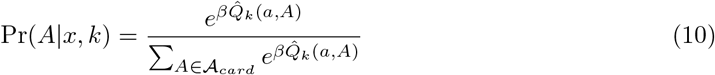

where *β* is an inverse temperature parameter that determines the tendency of the agents to exploit the highest estimated valued actions or to explore. Lower level primitive actions (needed to obtain the desired cardinal movement) were sampled using the mapping function:

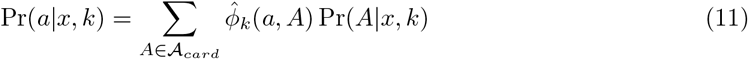

We first simulated the independent and joint clustering agents as well as the flat agent on 150 random task domains using the parameter values *γ* = 0.75, *β* = 5.0 and *α* = 1.0 (below we consider a more general parameter-independent analysis). Each of the four contexts was repeated 4 times for a total of 16 trials. The independent clustering agent completed the task more quickly than either other agent, completing all trials in an average of 205.2 (s=20.2) steps in comparison to 267.4 (s=22.4) and 263.5 (s=17.4) steps for the joint clustering and flat agents, respectively [Fig 2, B]. (We confirmed here and elsewhere that these differences were highly significant (e.g., here, the relevant comparisons are a minimum of *p* < 1e—77)). Repeating these simulations with agents required to estimate the full transition function (instead of just the mapping function) led to the same pattern of results, with the independent clustering agent completing the tasks in fewer steps than either the joint clustering or flat agents [Fig S1, A].

In this case, the performance advantage of independent clustering is largely driven by faster learning of the reward function, as indexed by the KL-divergence between the agents’ estimates of the reward function compared to a flate learner [Fig 2, C, left]. In contrast, both joint and independent clustering show generalization benefit when learning the mapping function [Fig 2, C, right]. This difference reflects an information asymmetry: in a new context, more information is available earlier to an agent about the mappings than the rewards, given that the latter are largely experienced when reaching a goal. (For example, in these environments, the first action in a novel context yields 2 bits of information about the mappings and an average of 0.07 bits of information about the rewards). As a consequence of this asymmetry, observing an element of a mapping can facilitate generalization of the rest of the mapping via the likelihood function, whereas observing unrewarded squares in the grid world tells the agent little about the location of rewarded squares.

In sum, as expected, independent clustering exhibited advantages over joint clustering in a task environment for which the transition and reward functions were orthogonally linked across contexts.

### Simulation 2: Dependence in Task Statistics

We next simulated all three agents on separate task domain in which there was a discoverable relationship between the reward and mapping functions across contexts, such that knowledge about one function is informative about the other. There were with four orthogonal reward functions and four orthogonal mappings across eight contexts, with each pairing of a reward and mapping function repeated across two contexts, permitting generalization [Figure 3, A]. As before, 150 random task domains were simulated for each model using the parameter values *γ* = 0.75, *β* = 5.0 and *α* = 1.0. Each of the eight contexts was repeated 4 times for a total of 32 trials. In these simulations, both clustering agents show a generalization benefit, completing the task more quickly than the flat agent [Fig 3, B]. The joint clustering agent showed the largest generalization benefit, completing all trials in average of 384.2 (s=23.6) steps in comparison to 441.5 (s=33.4) for the independent clustering agent and 526.0 (s=26.4) steps for the flat agents. Again, these differences were highly significant and the agents that estimated the full transition function displayed the same pattern of results [Fig S1, B].

**Fig 3.**
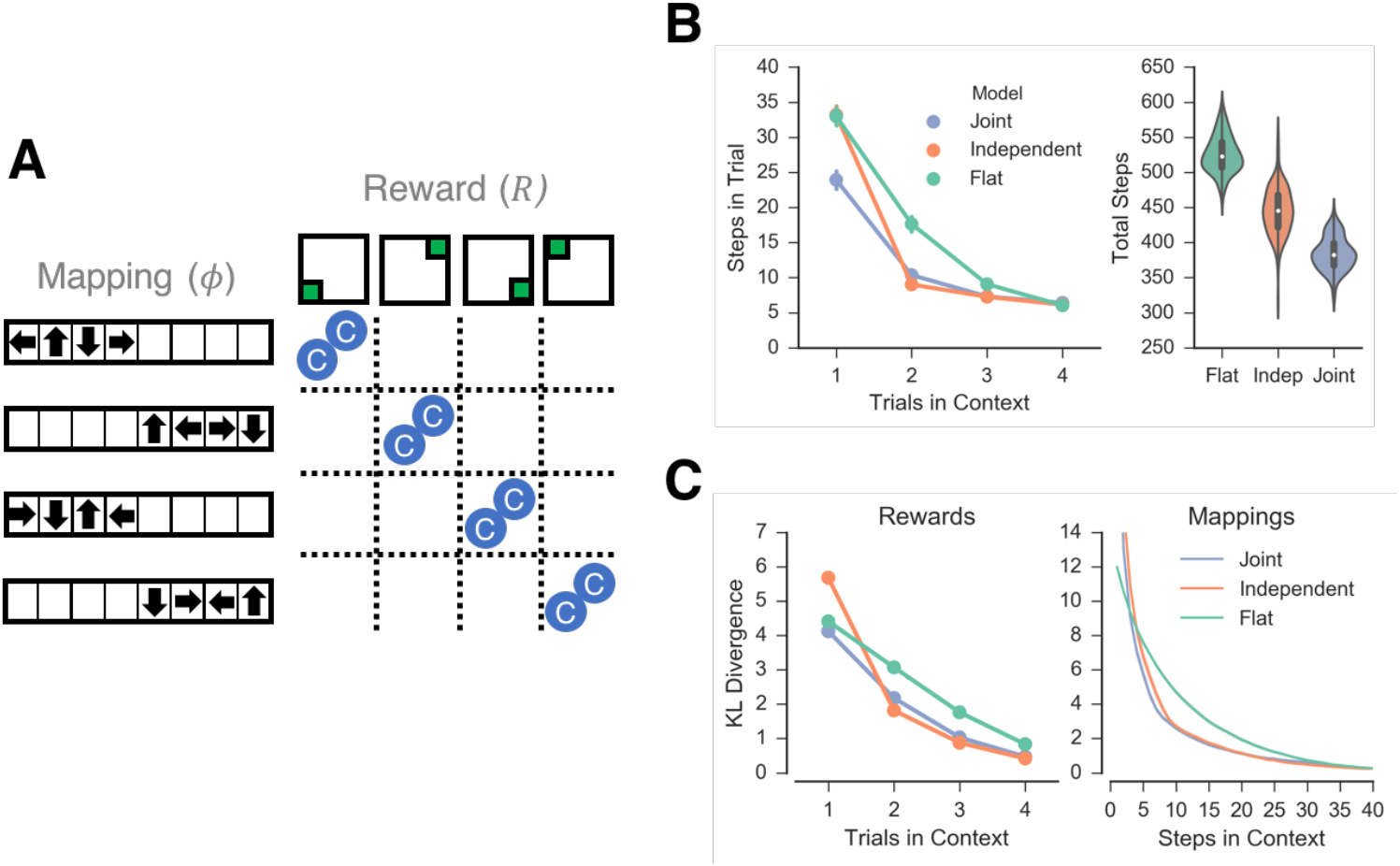
Simulation 2. A: Schematic representation of the second task domain. Eight contexts (blue circles) were simulated, each paired with a combination of one of four orthogonal reward functions and one of four mappings, such that each pairing was repeated across two contexts, providing a discoverable relationship. B: Number of steps taken by each agent shown across trials within a single context (left) and over all trials (right). C: KL-divergence of the models’ estimates of the reward (left) and mapping (right) functions as a function of time.

As is the previous simulations, the differences in performance between the clustering agents was largely driven by learning of the reward function. Both the independent and joint clustering agents had similar estimates of mapping functions across time [Fig 3, C, right] whereas the independent clustering agent uniquely shows an initial deficit in generalization of the reward function, as measured by KL divergence in the first trial in a context [Fig 3, C, left]. The difference in performance between the two clustering models largely occurs for the first trial in a new in a new context, during which time the joint clustering agent had a better estimate of the reward function. As before, this reflects an information asymmetry between mappings and rewards.

### Simulation 3: Compounding effects in the “diabolical rooms” problem

Generalization is almost synonymous with a reduction of exploration costs: if an agent generalizes effectively, it can determine a suitable policy without fully exploring a new context. In the above simulations, exploration costs were uniform across all contexts. But in real world situations, the cost of exploring can compound as a person progresses through a task. Exploration can become more costly as resources get scarce: for example, on a long drive it is far more costly to drive around looking for a gas station with an empty tank than with a full one because running out of gas is more likely. Likewise, in a video game where taking a wrong action can mean starting over, it is more costly for an RL agent to randomly explore near the end of the game than at the beginning. Thus, the benefits and costs of generalization can compound in task with a sequential structure over multiple subgoals in ways that are not often apparent in a more restricted task domain. Here, we consider a set of task domains in which each context has a different exploration cost, which increases across time.

We define a modified ‘rooms’ problem as a task domain in which an agent has to navigate a series of rooms (individual grid worlds) to reach a goal location in the last room [Fig 4, A]. In each room, the agent must choose one of three doors, one of which will advance the agent to the next room, whereas the other two doors will return the agent to the starting position of the very first room (hence the ‘diabolical’ descriptor). Additionally, the mappings that link actions to cardinal movements can vary from room to room, such that the agent has to discover this mapping function separately from the location of the reward (goal door). All of the rooms are visited in order such that if an agent chooses a door that returns it to the starting location, it will need to visit each room before it can explore a new door. Consequently, the cost of exploring a door in a new room increases with each newly encountered room.

**Fig 4.**
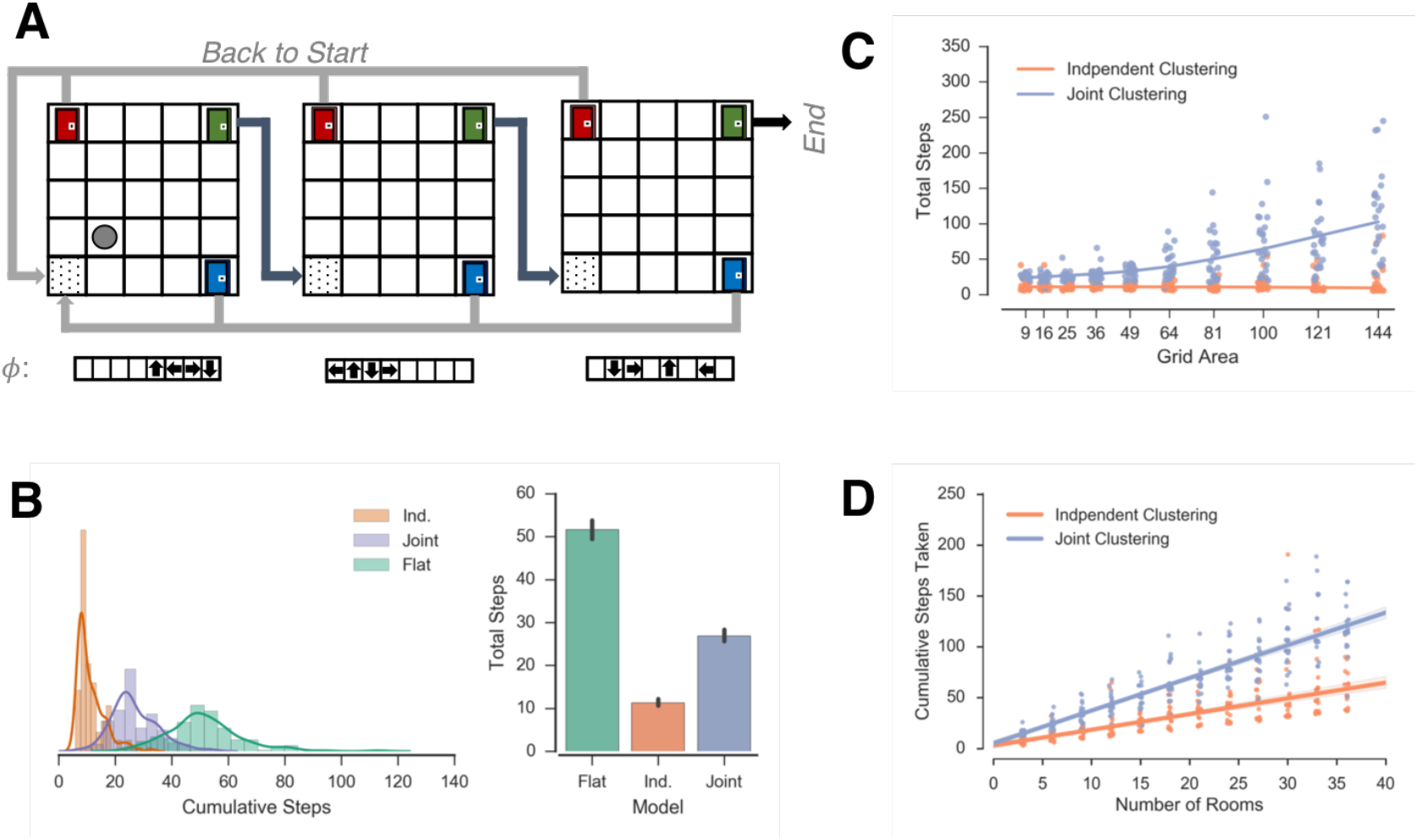
“Diabolic Rooms Problem”. *A*: Schamatic diagram of rooms problem. Agents enter a room and choose a door to navigate to the next room. Choosing the correct door (green) leads to the next room while choosing the other two doors leads to the start of the task. The agent learns three mappings across rooms *B*: Distribution of steps taken to solve the task by the three agents (left) and median of the distributions (right). *C*,*D*: Regression of the number of steps to complete the task as a function of grid area (C) and the number of rooms in the task (D) for the joint and independent clustering agents.

Botvinick et al. [21] have previously used the original ‘rooms’ problem introduced by Sutton et al. [20] to motivate the benefit of the “options” hierarchical RL framework as a method of reducing computational steps. However, in the traditional options framework, there is no method for reusing options across different parts of the state-space (for example, from room to room). Each option needs to be independently defined for the portion of the state-space it covers. In contrast, hierarchical clustering agents that decompose the state space can facilitate generalization in the rooms problem by reusing task structures when appropriate [3]. However, because it was a joint clustering agent, this previous work would not allow for separate re-use of mapping and reward functions.

In this new, diabolical, variant of the rooms problem, we have afforded the opportunity for reuse of subgoals across rooms, but have modified the task not only to allow for different mapping functions, but where there is a large cost when the appropriately learned subgoal (choosing the correct door) is not reused, and where this cost is varied parametrically by changing the size or number of the rooms. The rooms problem here is qualitatively similar to the “RAM combination-lock environments” used by Leffler and colleagues [9] to show that organizing states into classes with reusable properties (analogous to clusters presented in the present work) can drastically reduce exploration costs. In the RAM combination-lock environments, agents navigated through a linear series of states, in which one action would take agents to the next state, another to a goal state, and all others back to the start. The rooms task environment presented here is highly similar but allows us to vary the cost of exploring each room parametrically by varying its size.

We simulated an independent clustering agent, a joint clustering agent, and a flat agent on a series of rooms problems with the parameters *α* = 1.0, *γ* = 0.80 and *β* = 5.0. Each room was represented as a new context (for example, to simulate differences in surface features). There were three doors in the corners of the room, and the same door advanced the agent to the next room for every room (for simplicity, but without loss of generality - i.e., the same conclusions apply if the rewarded door would change across contexts). Agents received a binary reward for selecting the door that advanced to the next room.

In the first set of simulations, we simulated the agents in 6 rooms, each comprising a 6×6 grid world, with three mappings *∅_c_* ∈ {*∅*_1_, *∅*_2_, *∅*_3_}, each repeated once. Because the cost of exploration compounds as an agent progresses through a task, the ordering of the rooms affects the exploration costs. For simplicity, we simulated a fixed order of the mappings encountered by rooms, defined by the sequence *X*^∅^ = *∅*_1_*∅*_1_*∅*_2_*∅*_2_*∅*_3_*∅*_3_. In this task domain, independent clustering performed the best with both clustering agents show a generalization benefit as compared to the flat agent [Fig 4, B], with the flat agent completing the task in approximately 1.9x and 4.5x more total steps than joint and independent clustering, respectively.

We further explored how these exploration costs change parametrically with the geometry of the environment. First, we varied the dimensions of each room from 3×3 to 12−12. While both clustering models show increased exploration costs as the area of the grid world increases, the exploration costs for the joint clustering model grow at a faster exponential rate than the independent clustering model [Fig 4, C]. Similarly, we can increase the exploration costs by increasing the number of rooms in the task domain. We varied the number of rooms in the rooms in the task domain from 3 to 27 in increments of three. As before, the same door in all rooms advanced the agent and three mappings were repeated across the rooms. The order of the mappings encountered is defined by the sequence 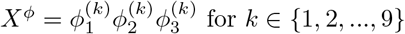, where *k* is the number of times a mapping is repeated. Again, both clustering agents experience an increased cost of exploration as the number of rooms increases, but the cost of exploration increases at a faster linear rate for the joint clustering agent than the independent clustering agent [Fig 4, D].

Thus, in environments where the benefits of generalization compound across time, difference between strategies can be dramatic. Here, we have simulated an environment in which independent clustering leads to better generalization than joint clustering but we could equivalently create an example in which joint clustering leads to better performance (for example, joint clustering would do better if each mapping uniquely predicted the correct door). Consequently, any fixed strategy has the potential to face an exploration costs that grows exponentially with the complexity of the task domain.

### Information Theoretic Analysis

Thus far, we have examined the performance of hierarchical clustering variants in specific situations in order to demonstrate a tradeoff between strategies. However, while these examples are illustrative, they impose strong assumptions about the task domain, the agents’ knowledge of its structure, exploration policies and planning. In contrast, we are more concerned with the suitability of generalization across ecological environments, rather than the specific task domains we have simulated and the assumptions of planning and exploration.

To make a more general normative claim, it is desirable to abstract away the implementation and strictly address the normative basis of context-popularity based clustering as a generalization algorithm by itself. While addressing optimality requires knowledge of the generative process of ecological environments, which is beyond the scope of the current work, we can more formally and generally assess when, and under what conditions, each of the clustering models might be more suitable than the other.

To do so, we can frame generalization as a classification problem and quantify how well an agent correctly identifies the cluster in which a context belongs without regard to learning the associated task statistics. This simplifying assumption allows us to examine the CRP as a mechanism for generalization and abstracts away the effect of the likelihood function on generalization. Let *k* ∈ *K* be a cluster associated with a Markov decision problem and let context *c* ∈ *C* be a context experienced by the agent. Given a history of experienced contexts and associated clusters {*c*_1:*n*_, *k*_1:*m*_}, we cast the problem of generalization as learning the classification function *k* = *f*(*c*) that minimizes the risk of misclassification.^1^

More formally, we define risk as the expectation E[*L*(*p*, *f*)], where *L*(*p*, *f*) is the loss function for misclassification and *p* is the generative distribution over new contexts *p* = Pr(*k*|*c*_t+1_). For our purposes here, we will abstract away domain specificity in the loss function *L*(*p*, *f*). Because the CRP is a probability distribution, a reasonable domain-independent loss function is the information gain between the CRP’s estimate of the probability of *k* and the realized outcome, or

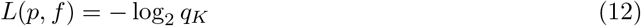

where *q_k_* is the CRP’s estimate of the probability of the observed cluster *k* in context *c*. The misclassification risk is thus the cross entropy between the CRP and the generative distribution:

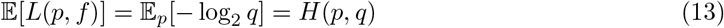

Thus, by casting generalization as classification and assuming information gain as a domain-general loss function, we are in effect evaluating the degree to which the CRP estimates the generative distribution. Risk is minimized when *q* = *p*, that is, when the CRP perfectly estimates the generative process. There is no upper bound to poor performance, but in many task domains it is possible to make a naïve guess over the space of clusters (for example, a uniform distribution over a known set of clusters). Because any useful generalization model will be better than a naïve guess, we can evaluate whether the CRP will lead to lower information gain than a naïve estimate in different task domains as a function of their statistics. We consider this more quantitatively in the appendix, but the result is intuitive: overall, the CRP will facilitate generalization when the generalization process is more predictable (less entropic) and the CRP can lead to worse than chance performance in sufficiently unpredictable domains [Fig S2].

#### Independent vs Joint clustering

A question of primary interest is under what conditions is it better to cluster aspects of the task structure (such as reward functions and transition or mapping functions) independently or jointly. To do so, we rely on the assumption that a reward function is typically substantially sparser than a transition function: as an agent interacts with a task it will gain more information about the transition function early in the task than it will about the reward function (unless the environment is very rich with rewards at most locations, in which case any random agent would perform well).

Consequently, for an agent that clusters rewards and transitions together, the information gained about transitions will dominate the likelihood function, such that the inference of rewards can be thought of as approximately conditional on knowledge of the transition structure. Conversely, an agent that clusters rewards and transition independently will not consider the mapping information when it predicts the reward function. Thus, we can compare the two agents by evaluating the consequences of clustering rewards conditional on transitions as compared to clustering rewards independent of transitions. Formally, we can consider the comparison of independent and joint clustering as the comparison between two different classifiers, *R* = *f*(*c*) and *R* = *f*(*c*, *T*), one of which classifies reward functions solely as a function of contexts and the other as a function of contexts and observed transition statistics.

Interestingly, this approximation leads to the conclusion that independent clustering is a simpler statistical model than joint clustering. Estimating a marginal distribution is a simpler statistical problem than estimating its composing set of conditional distributions. As such, we might expect a tradeoff where independent clustering provides better generalization with little experience whereas joint clustering provides better generalization asymptotically. We can evaluate the latter claim by noting that given random variables *R* and *T*, *H*(*R*|*T*) ≤ *H*(*R*). This statement implies that given a known joint distribution between two random variables, knowledge of one of the random variables cannot increase the uncertainty of the other; an agent can simply learn when there is no relation between the two, in which case the joint distribution doesn’t hurt. Intuitively, this claim is based on the notion that more information is always better (or at least, no worse) in the long run.

Note that this relationship is only guaranteed if the true generative process is known and as such, experience in the task domain plays an important role. As we discuss in the following section, there needs to be sufficient experience to determine whether the joint distribution is useful or not (and in fact, assuming conditional dependence when there is no such relationship can slow down learning dramatically). Nonetheless, the CRP prior will converge asymptotically on the conditional and marginal distributions, Pr(*R*|*T*) and Pr(*R*), for the joint and independent clustering agents, respectively. If we consider the CRP to be an estimator, we can define its bias as

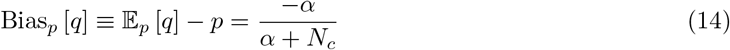

where *N_c_* is the total number of contexts observed and *α* is the concentration parameter governing new clusters. Asymptotically, the CRP is unbiased as lim_*N*_*c*_→∞1_ Bias_*p*_[*q*] = 0 and the CRP converges to the generative distribution. As a consequence, joint clustering has lower information gain than independent clustering asymptotically as the CRP for a joint clustering agent will converge to the conditional distribution Pr(*R*|*T*) whereas the CRP for an independent clustering agent will converge to the marginal distribution Pr(*R*).

#### Mutual Information

As alluded to above, that joint clustering is guaranteed to produce a better estimate is only true as *N_c_* → ∞ 1 whereas here we are concerned with task domains in which an agent has little experience. Intuitively, we might expect independent clustering to be favorable in task domains where there is no relationship between transitions and rewards. Conversely, we might expect joint clustering to be more favorable when there is relationship between transition and rewards. While we considered two extremes of these cases in simulations 1 and 2, we can also vary the relationship parametrically. Formally, we can consider the relationship between transitions and rewards with mutual information. Let each context *c* be associated with a reward function *R* ∈ *R* and a transition function *T* ∈ *T* and let Pr(*R*, *T*) be the joint distribution of *R* and *T* in unobserved contexts. The mutual information between *R* and *T* is defined

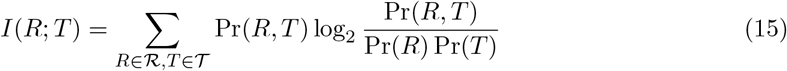

Mutual information represents the degree to which knowledge of either variable reduces the uncertainty (entropy) of the other, and can be used as a metric of the degree to which a task environment should be considered independent or not. As such, it satisfies 0 ≤ *I*(*R*;*T*) ≤ min (*H*(*R*),*H*(*T*))

To evaluate how mutual information affects the relative performance of independent and joint clustering, we constructed a series of task domains that allow us to monotonically increase *I*(*R*; *T*) by with a single parameter *m* while holding all else constant. We define *R* = {*A*,*B*} and *T* = {1, 2} and we define two sequences *X^R^* and *X^T^* such that *X*_i_^R^ and *X*_i_^T^ are the reward and transition function for context *c_i_*. We define the sequence *X^R^* = *A*^(2n)^*B*^(2n)^, where *A*^(k)^ refers to a *k* repeats of *A*, and the sequence *X^T^* = 1^(*n*+*m*)^2^(2*n*)^1^(*n*-*m*)^, where 1^(*k*)^ refers to *k* repeats of 1. To provide a concrete example, if *n* = 2 and *m* = 0, the sequence *X^R^* = *AAAABBBB* and the sequence *X^T^* = 11222211. Similarity, if *n* = 2 and *m* = 2, then the sequence *X^R^* is unchanged while *X^T^* is now 11112222. Critically, these sequences have the property that *I*(*R*;*T*) = 0 if and only if *m* = 0 and that *I*(*R*; *T*) monotonically increases with *m*. In other words, the residual uncertainty of *R* given *T*, *H*(*R*|*T*), declines as a function of *m*. This allows us to vary *I*(*R*; *T*) independently of all other factors by changing the value of *m*.

We evaluated the relative performance of the independent and joint clustering agents by using the CRP to predict the sequence *X^R^*, either independent of *X^T^* (modeling independent clustering), or conditionally dependent on *X^T^* (modeling joint clustering) for values of *n* = 5 and *m* = [0, 5]. For these simulations, we first assume both sequences *X^R^* and *X^T^* are noiseless, but below we show that noise parametrically affects these conclusions. For low values of *m* (*m* ≤ 2), independent clustering provides a larger generalization benefit [Fig 5, left]. This has the intuitive explanation that as the features of the domain becomes more independent, independent clustering provides better generalization. Importantly, there are cases in which independent clustering provides a better evidence even thought there is non-zero mutual information [0 < *I*(*R*; *T*) ≲ 0.2bits].

**Fig 5.**
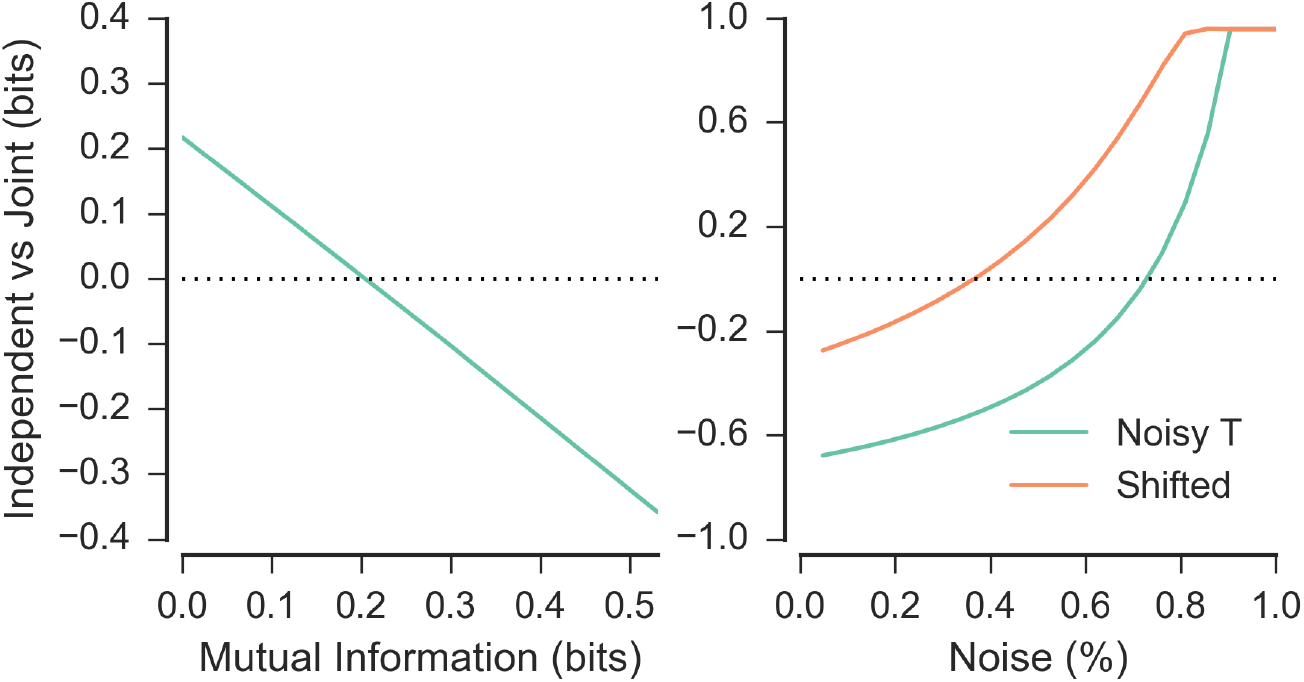
Performance of independent vs. joint clustering in predicting a sequence *X_R_*, measured in bits of information gained by observation of each item. *Left*: Relative performance of independent clustering over joint clustering as a function of mutual information between the rewards and transitions. *Right*: Noise in observation of *X_T_* sequences parametrically increases advantage for independent clustering. Green line shows relative performance in sequences with no residual uncertainty in *R* given *T* (perfect correspondence), orange line shows relative performance for a sequence with residual uncertainty *H*(*R*|*T*) > 0bits.

#### Noise in observation of transition functions

Above, we assumed that transition functions were fully observable with no uncertainty. But in many real-world scenarios, the relevant state variables are only partially observable and the transition functions may be stochastic. In this section, we therefore relax this noiseless assumption to characterize the effect of noisy observations on inference and generalization. As before, we model generalization as the degree of predictability of 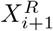 given 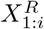 either independent of *X^T^* (independent clustering) or conditionally on *X^T^* (joint clustering).

We first construct reward and transition sequences in which knowledge of the transition function completely reduces the uncertainty about the reward function (*I*(*R;T*) *= H*(*R*), and hence *H*(*R|T*) *=* 0). Consider **X^R^* = A*^(20)^*BCD* and *X^T^* = 1(^20^)234, where *A*^(20)^ and 1(^20^) refer to 20 repeats of *A* and 1, respectively. As such, we would expect joint clustering to produce better generalization than independent clustering if observations are noiseless. To simulate noise / partial observability we assume each observation of 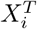 is mis-identified as some new function *T*\* ∉ {1,2,3,4} with some probability. Importantly, this simulation has the desideratum that noise does not affect *I*(*R;T*) or *H*(*R*) themselves (the true generative functions). We compare the inference of **X^R^** independent of *X^T^* (modeling independent clustering) or conditionally dependent on *X^T^* (modeling joint clustering) for various noise levels *σ* = [0, 1.0]. When noise is sufficiently high (*σ* > 0.71) independent clustering produces a better estimate of **X^R^** than joint clustering [Fig 5, right, green line] even for this extreme case where the two functions have perfect mutual information.

Next, we assessed how mutual information and noise interact by decreasing the correspondence of the sequences. As noted above, in the sequences used above (**X^R^* = A*^(20)^ *BCD* and *X^T^* = 1^(20)^234), there is no residual uncertainty of *R* given *T*. We can decrease the correspondence between the two sequences by shifting *X^T^* by one to 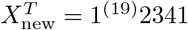 and thus decrease the mutual information 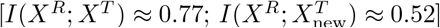. As we have previously noted, this decline in mutual information reduces the benefit of joint clustering, *ceteris paribus*. Nonetheless there is still a strong relationship between rewards and transition that can be leveraged by joint clustering.

Simulating independent and joint clustering on these new sequences as a function of *σ* = [0 1.0] reveals a lower level of noise needed to see a benefit of independent clustering [Fig 5, right]. As expected joint clustering provides a better estimate in the no-noise case, as well as for low noise levels (*σ* < 0.33) as it can take advantage of the shared structure, while independent clustering results in a better estimate for larger noise levels. Importantly, these effect are cumulative; in ecological settings where observations are noisy and there is only weak mutual information, independent clustering will likely provide a better estimate of the prior over rewards.

### Meta-agent

Given that the optimality of each fixed strategy varies as a function of the statistics of the task domain, a natural question is whether a single agent could optimize its choice of strategy effectively by somehow tracking those statistics. In other words, can an agent infer whether the overall statistics are more indicative of a joint or independent structure and capitalize accordingly? Here, we address this question by implementing a meta-agent that infers the correct policy across the two strategies (below we also consider a simple model-free RL heuristic for arbitrating between agents, which produces qualitatively similar results).

For any given fixed strategy, the optimal policy maximizes the expected discounted future rewards and is defined by equation 1. Let 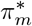 be the optimal policy for model *m*. We are interested in whether 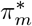 is the global optimal policy π^*^, which we can define probabilistically as

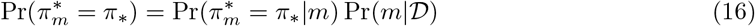

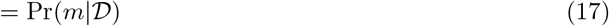

where Pr(*m*|*D*) is the Bayesian model evidence and where Pr(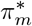 = π_*_|*m*) ≡ 1. The Bayesian model evidence is, as usual,

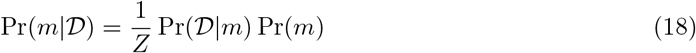

where Pr(*D*|*m*) is the likelihood of observations under the model, Pr(*m*) is the prior over models and Z is the normalizing constant. The likelihood function for the independent and joint clustering models is the product of the mapping and reward functions (as defined conditionally for a cluster in equations 6, 7 and 8). However, estimates of the mapping function is highly similar for both models (Fig. 2C, Fig. 3C). Thus, as a simplifying assumption, we can approximate the model evidence with how well each model predicts reward:

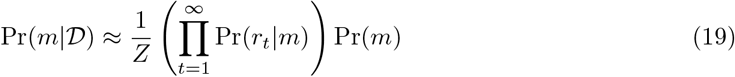

where *r_t_* is the reward collected at time *t*. Independent and joint clustering can be interpreted as special cases of this meta-learning agent with the strong prior Pr(*m*) = 1. Under a uniform prior over the models, this strategy reduces to choosing the agent based on how well it predicts reward, an approach repeatedly used in meta-learning agents [3,23,24].

We simulated the meta-learning agent with a uniform prior over the two models and used Thompson sampling [25] to sample a policy from joint and independent clustering at the beginning of each trial [Fig 6, A]. In the first task domain, where independent clustering results in better performance than joint clustering, the performance of the meta-agent more closely matched the performance of the independent clustering agent [Fig 6, B]. The meta-agent completed the task in an average of 235.2 (s=35.3) steps compared to 205.2 (s=20.2) and 267.5 (s=22.4) steps for the independent and joint clustering agents. In addition, the meta-agent became more likely to choose the policy of the independent clustering agent over time [Fig 6D]. In the second task domain, where joint clustering outperformed independent clustering, the meta-agent completed the task in an average of 417.5 (s=42.0) steps compared to an average of 384.2 (s=21.2) and 441.6 (s=35.9) steps for the joint and independent clustering agents, respectively. Likewise, overtime the meta-agent was more likely to choose the policy of the joint clustering agent [Fig 6, E].

**Fig 6.**
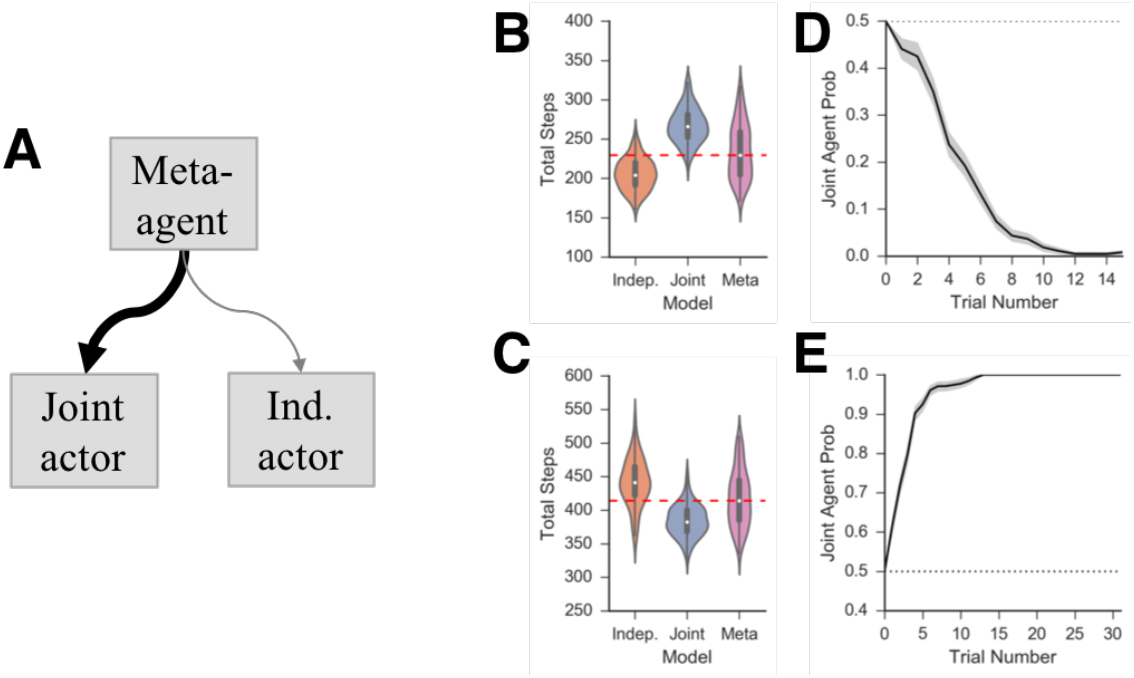
Meta-Agent. *A*: On each trial, the meta-agent samples the policy of joint or independent actor based on model evidence for each strategy. Both agents, and their model evidences, are updated at each time step. *B*: Overall performance of independent, joint and meta agents on simulation 1 *C*: Overall performance of independent, joint and meta agents on simulation 2 *D,E*: Probability of the selecting the policy joint clustering over time in simulation 1 (*D*) and simulation 2 (*E*).

A computationally simple approximation to estimating the model responsibilities is to select agents as a function of their estimated value. In this approximation, a reward prediction error learning rule estimates the value for each model, *Q_m_*, according to the updating rule:

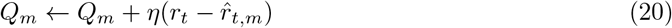

where *η* is a learning rate and 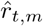 is the reward predicted by the model at time *t*. These values can be used to sample the models via a softmax decision rule

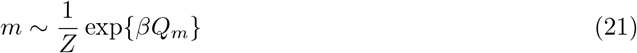

Simulation with this arbitration strategy with the parameter values *β* = 5.0, *η* = 0.2 led to a qualitatively similar pattern of results. Performance in both simulations 1 and 2 were not distinguishable from the Bayesian meta agent, with the agent completing simulation 1 in 237.2 (s=41.2) steps as compared to the 235.2 (s=35.3) found for the Bayesian implementation (*p* < 0.65) and completing simulation 2 in 418.7 (s=45.9) steps as compared to 417.5 (s=42.0) steps for the Bayesian agent (*p* < .81).

Thus, while for both inference and RL versions, the meta-agent did not equal the performance of the best agent in either environment, it outperformed the worse of the two agents in both environments. Normatively, this is a useful property if an agent cares about minimizing the worst possible outcome across unknown task domains (as opposed to maximizing their performance within a single domain), similar to a minimax decision rule in decision theory [26]. This can be advantageous if agent has little information about the distribution of task domains and if the costs of choosing the wrong strategy are large as in the ‘diabolical rooms’ problem. Furthermore, while we have used a uniform prior over the two strategies, varying the prior may result in a better strategy for a given set of task domains.

In our information theoretic analysis above, we showed that the task statistics determines the normative strategy depending on which agent is more efficacious in reducing Bayesian surprise about reward. The meta-learning agent capitalizes on this same intuition by using predicted rewards to arbitrate among strategies. More specifically, we argued that the normative value of each strategy varies with mutual information between rewards and mappings. Thus, we assessed whether the the meta-learning agent is also sensitive to mutual information, without calculating it directly, and hence be more likely to choose joint clustering when the mutual information is higher.

In simulations 1 and 2 above, we calculated the mutual information each time a new context was added and used this to predict the probability of selecting the joint agent at the end of that trial.

Specifically, we define

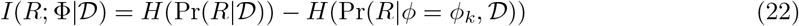

where Pr(*R*|*D*)) is the probability each location is rewarded across all contexts seen so far and Pr(*R*|∅ = *∅_k_*, *D*)) is conditioned on the current mapping. Using logistic regression, we find that *I*(*R*; Φ|*D*) positively correlates with the probability of selecting the joint agent across both simulations, consistent with expectations [Simulation 1: *β* = 3.3, *p* < 0.003; Simulation 2: *β* = 9.4, *p* < 5 × 10^−41^].

## Discussion

In this paper, we provide two alternative models of context-based clustering for the purpose of generalization, a *joint clustering* agent generalizes reward and transition functions together and an *independent clustering* agent that separately generalizes reward and transition functions. These models are motivated by human learning and performance, which is thought to be structured and compositional [27–29].

Generalization can be seen as a solution to a dimensionality problem. In real-world problems, perceptual space is typically high-dimensional. In order to learn a policy, agents need to learn a mapping between the high-dimensional perceptual space and the effector space. Learning this mapping can require a large set of training data, perhaps much larger than a human would have access to [12,21]. Clustering can reduce the dimensionality by projecting the perceptual space onto a lower dimensional latent space in which multiple percepts share the same latent representation. Thus, an agent does not need to learn a policy over the full state space but over the lower dimensional latent space. This is an explicit assumption of the models presented here as well as in other clustering models of human generalization, allowing agents to collapse across irrelevant features and preventing interference between stimulus-response mappings across latent distinct rules [3,23]. Related principles have been explored in lifelong learning [6–8], object-based, and symbol-based approaches [11,30–33].

Incorporating compositionality takes this argument further, as multiple policies often share component features. For example, playing the saxophone involves the use of the same movements to produce the same notes for different effect in different songs. Learning a policy as a direct mapping from the low level effector space to reward values fails to take advantage of the structure, even if that policy can be reused as a whole with another instrument. Thus, learning at the component level as opposed to the policy level reduces a high-dimensional problem into multiple lower-dimensional problems. While this adds the additional complexity of the choice of a good set of component features, here we argue the Markov decision process provides a natural decomposition into reward and transition functions. Importantly, this decomposition of the task structure is not equivalent to a decomposition of the policy, which is itself dependent on the joint reward and transition functions. Of course, other decompositions are also possible and useful (and not mutually exclusive). For example, the state-outcome and action-dependent state transition functions of the active inference framework can both be decomposed into “what” and “where” aspects [34,35]. While these functions, analogous to reward and transition functions, are linked by a shared latent state representation, this decomposition facilitate generalization across states that share features.

Regardless of the choice of component features, a compositional generalization model needs to make assumptions about the relationship between components. We argue here that the proper choice depends on the generative structure, which as an empirical matter, is largely unknown for the ecological environments faced by humans and artificial agents. As we demonstrated in the grid-world simulations above, when there is a strong relationship between components, an agent that assumes as much outperforms an agent that assumes no relationship, and vice-versa [Figs. 2 & 3]. With sufficient (and stationary) experience, we might expect a model that assumes a learnable relationship between components (joint clustering) to perform better in new contexts, since assuming a potential relationship between goals and mappings can be no worse asymptotically than assuming independence (i.e., the agent can simply learn that the correlation is zero). Nonetheless, how much experience is sufficient for joint clustering to provide a better model is difficult to define in general, and will depend on the statistics of the relationships and the combinatorial explosion of the state space that arises. Furthermore, noise or partial observability further complicates the picture: even when there is exploitable mutual information, independent clustering can yield a better estimate when experience is limited [Fig 5].

Why is this the case? It may appear puzzling given the asymptotic assurances that joint clustering will be no worse in stationary environments. Here, the comparison between classification and generalization is instructive. We can think of joint clustering in terms of estimating a joint distribution of the generative process and independent clustering in terms of estimating the marginal distribution for each component independently (similar to a naïve Bayes classifier). In this interpretation, independent clustering trades off an asymptotically worse estimate of the generative process for lower variance, with a bias equal to the mutual information between mappings and goals. In problems with limited experience, such as the type presented here, a biased classifier will often perform better than an asymptotically more accurate estimator because misclassification risk is more sensitive to variance than bias [36]. Thus, by ignoring the correlation structure and increasing the bias to generalize goals that are most popular overall, independent clustering may minimize its overall loss.

Intuitively, we can think of joint clustering as being potentially overly sensitive to noise. While over infinite time it is always better to estimate the correlations between components, in practice it may not be worth the cost of doing so. This happens when the correlations between transitions and rewards is weak or difficult to determine. For a human learner, an example might be the relationship between how hungry a person is and how heavy it is to carry a plate of food from a buffet. Learning this relationship is guaranteed to be asymptotically better than ignoring it but given the triviality of the benefit and the frequency of the context, it probably isn’t worth the exploration cost.

Previous models of compositional generalization have attempted to decompose the space of policies, rather than task structure, in to reusable pieces that can be re-executed [34, 37]. Because learned policies depend on both the reward and transition functions of a specific task, this decomposition implicitly generalizes these two sources of information together, and thus does not address the set of issues considered here (i.e., when the transition function is independent of the reward function across contexts). The same issue applies to the options framework and other hierarchical task representations [10,32,38–40]. As a consequence, reusing policy components will cause the agent to explore regions of the state space that have had high reward value in other contexts, which as we have shown may or may not be an adaptive strategy. For example, successful generalization in the “diabolical rooms” problem presented here, and the “finger sailing” task presented by Fermin and colleagues [14,15], requires a separation of reward from movement statistics. Indeed, the generalization of policy-dependent successor state representations works well only under small deviations of the reward or transition function [10,38,39]. Thus, the choice of components should be influenced by the robustness to changes in the reward and transition function, which will not necessarily linked to an individual policy.

From the perspective of human cognition, compositional representation provides the flexibility to create novel policies in a novel domains in a rule-governed manner. This flexibility, also known as *systematicity* or *generativity*, has long been thought to be a key feature of cognition [29, 41]. As Lake and colleagues note, a person can re-use the learned knowledge of the structure of a task to accomplish an arbitrary number of goals, such as winning a specific number of points in a video game [12]. Strongly linking component properties may impede the potential for systematicity by limiting the flexibility to recombine knowledge. As we have argued above, recombining reward and transition information may be particularly valuable, such that agents that can only generalize policies and reward-sensitive policy components may lack systematicity.

An altogether different possibility is that a mix of strategies is appropriate. While we argue that independent clustering is a simpler statistical problem than joint clustering, there are clearly cases where joint clustering is advantageous. As noted before, generalization of successor-state representation partially links transitions and rewards and is nonetheless sufficiently flexible to handle small deviations in policy [10, 38, 39]. Furthermore, joint clustering support simpler algorithms, such as the form of temporal difference learning algorithms thought to underlie human fronto-striatal learning [3, 42] as well as the proposed successor state representations recently proposed to underlie hippocampal-based planning [38,39,43]. As we have suggested with the meta-learning agent, trading off between joint and independent clustering can reduce the risk of the decision problem.

Furthermore, it is not known what a human learner would typically consider to constitute a higher order context variable separate from lower order state variables [3, 42]. Ecologically, the number of contexts a human could potentially encounter is quite high, in which case they would be able to form a more accurate estimate of the correlation structure between components over time. If this speculation is true, then one potential adaptive strategy would be to assume a weak relationship between components early in learning and increasingly relying on the correlation structure as the evidence supports it.

Thus, a hybrid system is supported by both computational and algorithmic considerations. From the perspective of biological implementation, the inference required for context-clustering based generalization can be approximated by a hierarchical cortico-basal ganglia learning system [3]. This framework could be extended to account for independent clustering by allowing for multiple cortical clusters separately representing reward and mapping functions, each of which is learnable by a neural network model [44]. Because joint clustering results in the same policy generalized to each context in a cluster, joint clustering does not require separately estimating the reward and transition functions and instead learned policies (such as stimulus-action values) can be generalized directly. This can obviate planning, a challenge for any biological model of any model-based control. Nonetheless, multiple lines of research suggest humans engage in model-based control [14,15,39,45,46] and human subjects can re-use arbitrary action-movement mappings (highly similar to the ones proposed here) for model-based control, suggesting a compositional representation potentially mediated by the dorsolateral prefrontal cortex, dorsomedial striatum and cerebellum [14,15].

Finally, while we have presented independent clustering as motivated by human capabilities for generalization, the question of whether human learning is better accounted for by independent or joint clustering, or a mixture of the two, remains an open question. While models are a generalization of previous models used to account for human behavior [2–4], they make separate testable predictions for human behavior. Joint clustering predicts that in a generalization task, human subjects will use transition formation to infer the location of an unknown goal. Independent clustering, in contrast, predicts human subjects will ignore transition information when searching for goals, and ignore goals when inferring the transition function. By providing humans subjects an initial set of contexts where the popularity of reward function varies across contexts as a function of the mapping, a novel set of test contexts can be chosen to differentiate the model predictions. Future work will address these predictions and the underlying brain mechanisms.

## Compositional clustering in task structure learning

### S1 Supporting Information

#### Grid world simulations with unknown transition functions

Here, replicate simulations 1 and 2 presented in the main article while relaxing the the assumptions the agent knows the spatial relationship between states. Instead, we consider the case in which an agent needs to learn the full transition functions over states, actions and successor states. The simulations presented here are the same as simulations 1 and 2 in the main article but we redefine the agents such that they no longer have access to the the transition function in terms of cardinal movements *T_c_*(*x,A,x*′).

We define the transition functions learned by the agents in terms of primitive actions *a* ∈ *A* as *f_c_*(*x, a, x*′) in context *c* by marginalizing over cardinal actions such that

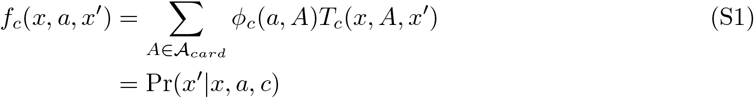

Correspondingly, the optimal policy can also be re-expressed in in terms of transition function *f_c_* as:

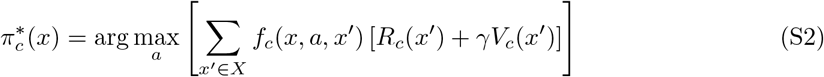

Likewise, the optimal value function is thus:

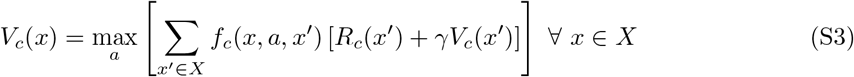

For the purpose of generalization, we assume the agents generalize the full transition functions, as opposed to generalizing the mapping functions presented in the main manuscript. As *T_c_*(*x, A, x*′) was previously assumed to be known, generalizing mappings can be seen as a specific case of generalizing full transition functions as a consequence of equation S1.

For the purpose of clustering contexts, the mapping function was used as a component of the likelihood function. Specifically, the likelihood function for the context-clustering assignments in joint clustering is the product of the mapping and reward functions *L*(*D*|*k*) = *∅_k_*(*a, A*)*R_k_*(*x, A*). Independent clustering assigns contexts into clusters separately for mappings and rewards, using *L*(*D*|*k*_∅_) = *∅_k_*(*a, A*) as the mapping cluster likelihood. As a consequence of equation S1, we can create the more general case of these two likelihood functions, respectively, with the following:

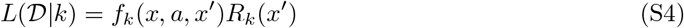

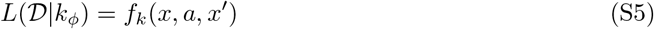

No further changes in the generative framework are needed to accommodate clustering full transition functions.

In the following simulations, agents estimated 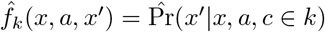 with maximum likelihood estimation, assuming independence between *f_k_*(*x, a, x*′) for all values of x and a. Action selection was performed by a combination of Thompson sampling [1] and ϵ-greedy exploration. For joint clustering, a single context-clustering hypothesis was sampled on each time step proportionally to its posterior probability. The estimates 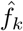 and 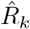 from sampled hypothesis were used to compute a state-action value function, defined

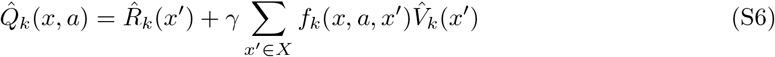

where the value function *V_k_*(*x*′) is generated from equation S3] via dynamic programming [2]. The sampled state-action value function is generated in the same manner for independent clustering, except that 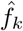 and 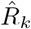 are sampled from the separate sets of clusters. Likewise, the flat model assumes the singular hypothesis that each context belongs to its own cluster and thus always samples the same hypothesis.

The state-action value function is used to generate a policy via an epsilon-greedy exploration rule where the action with the highest value 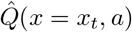 was chosen with probability 1 — ϵ and a random action was chosen with probability *ϵ* (ties were broken with equal probability).

##### Simulation 1

We first simulated the three agents on the same 150 random tasks presented in Simulation 1 of the manuscript. In this set of simulations, each of two reward functions and each of two transition functions were repeated across four contexts such that each context had a unique combination of reward and transition functions. The models were simulated using the parameter values *γ* = 0.75, *ϵ* = 0.5 and *α* = 1.0. As in the main manuscript, the independent clustering model learned the task more quickly than either of the other two models (*p* < 0.003 vs. joint; *p <* 0.004 vs. flat), completing all trials in an average of 1021.0 steps (s=377.2) in comparison to 1160.3 (s=408.3) and 1159.7 (s=424.9) steps for the the joint clustering and flat agents, respectively [Figure S1A].

##### Simulation 2

We then simulated the three agents on the same 150 random tasks presented in Simulation 2 of the manuscript. In this set of simulations, each of four reward functions and each of four transition functions were repeated across eight contexts such that each paring of reward and transition functions was repeated across two contexts. As in the main manuscript, the joint clustering model learned the task more quickly than either of the other two models (*p* < 0.08 vs. independent; *p* < 10^−26^ vs. flat), completing all trials in an average of 1485.8 steps (s=496.4) in comparison to 1583.6 (s=449.1) and 2820.0 (s=653.8) steps for the the independent clustering and flat agents, respectively [Figure S1B].

---

**Supplementary Figure 1.**
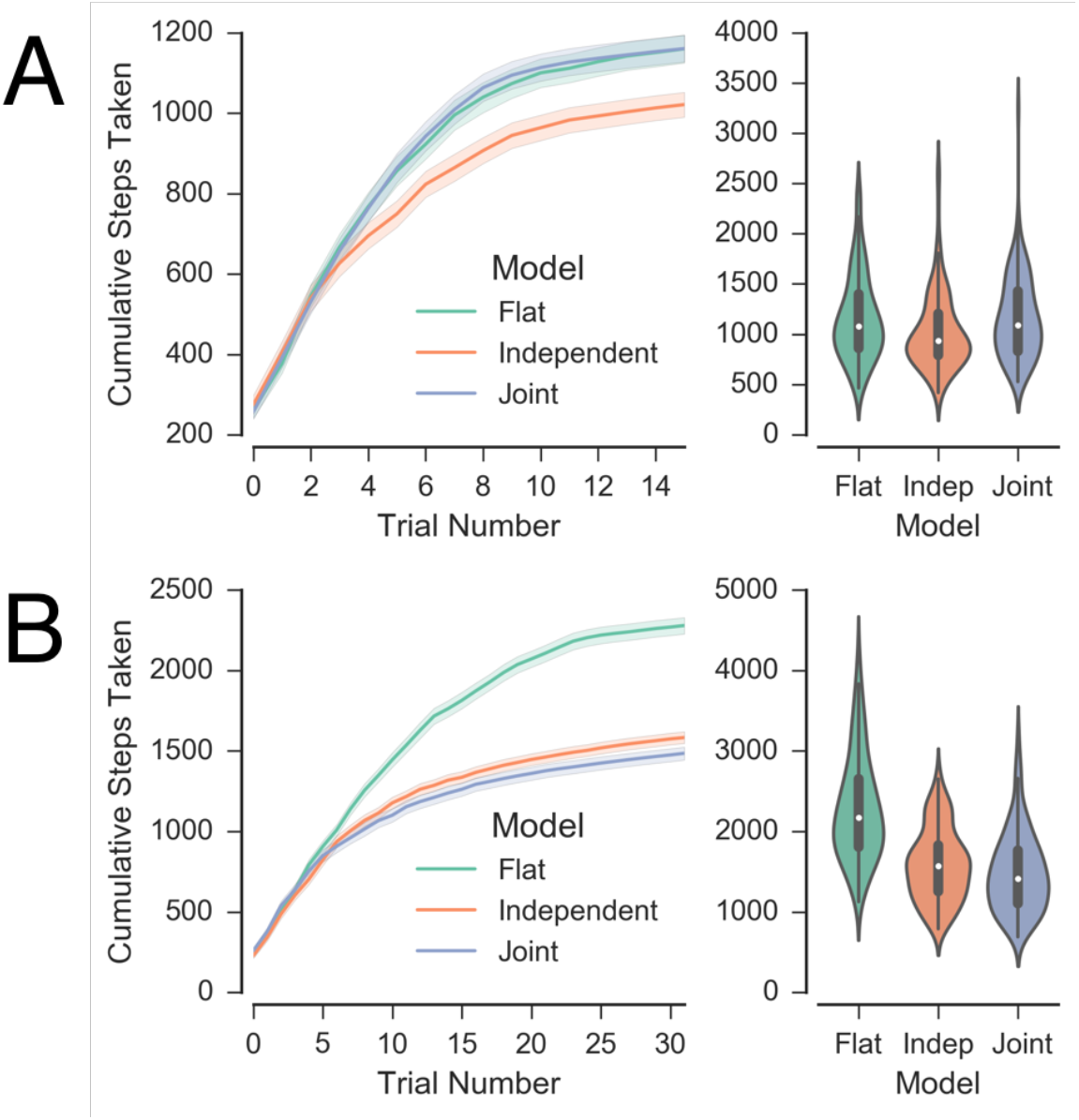
A: Agents’ performance learning full transition function in Simulation 1. Left: Cumulative number of steps taken by each model as a function of trials. Fewer steps represents better performance. Right: Distribution of total number of steps required to complete the task for each agent. B: Agents’ performance function in Simulation 2.

## Compositional clustering in task structure learning

### S2 Supporting Information

#### CRP performance as a function of task structure

One question we can ask is how well the CRP prior generalizes as a function of the underlying structure of the task domain. Intuitively, we might expect that a generalization agent should show a generalization benefit in highly structured task domains and show a less of a generalization benefit in unstructured domains. We can thus define a set of restricted task domains that vary in their structure. Specifically, we can vary the predictability of the generative process *p* and evaluate the agent as a function of this predictability.

Let *K* = {*A, B, C, D*} be the set of possible clusters in a task domain with probabilities *P* = {*p_A_,p_B_,p_C_,p_D_*}. Let *X* = *ABCD* represent a sequence of contexts and cluster identities experienced by an agent, such that cluster *A* is experienced in context *c*_1_, B is experienced in context *c*_2_, etc. We assume that the cluster identity is observable from the statistics of the associated MDP and that the agent knows the members of the set *K*.

We want to evaluate the ability of the CRP prior to predict each cluster in the sequence, conditional on its own history. We do so by calculating the expected loss experienced by the CRP over the sequence *X*. We define our loss function over sequence *X* as

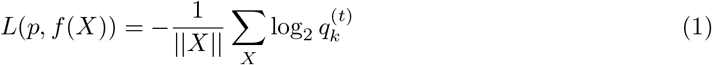

where 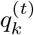 is the CRP’s probability estimate for the value *k* = *X*_t_ given *X*_1:*t*-1_. As noted above, this loss function is equivalent to the cross entropy *H*(*q, p*) between the CRP and the generative process. That is, *H*(*q, p*) is the average degree of unpredictability (in bits of information content) of experiencing each MDP given the estimate *q*. We can assess *H*(*q,p*) for the CRP by updating the predictive distribution in each context and probing its estimate for the subsequent context. As the CRP is exchangeable [1], *H*(*q, p*) is invariant to the order of the sequence, though the MAP approximation in the previous set of simulations can introduce order effects.

We can similarly quantify the degree of predictability of *X* by evaluating the entropy of the sequence, defined 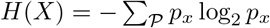. Here, we define a sequence *X*^(*n*)^ to allow us to monotonically decrease H(X) with n. Let *X*^(*n*)^ be the sequence *A*^(*n*)^*BCD*, where *n* denotes the number of times A appears in the sequence. For example, *X*^(1)^ is the sequence ABCD and *X*^(3)^ is the sequence AAABCD. For simplicity, we assume the probability distribution *P* over the ensemble *K* is exchangeable and that the probabilities over the members of its ensemble are proportional to their frequency in the sequence such that *p*_k_ = *N*_k_/||*X*|| where *N*_*k*_ is the number of times *k* appears in sequence *X*. Consequently, the entropy of the sequence *X*^(1)^ is *H*(*X*^(1)^) = 2 bits and the entropy of the sequence *X*^(3^ is *H*(*X*^(3)^) ≈ 1.79bits. As *n* approaches infinity, the entropy *H*(*X*^(*n*)^) asymptotically approaches 0 bits. Intuitively, as *A* is repeated more often in the sequence, the sequence is more predictable (lower entropy). It is important to note that the sequence predictability does not depend on order. Because the CRP is exchangeable, it will have the same predictive error for the sequences ABCDABCD and ADBCDBAC. Order-dependent predictability is beyond of the scope of the current work.

We evaluated the CRP on *X*^(*n*)^ for *n* = [1,100] and compared it to a naïve guess (uniform distribution over M). Because the CRP is parameterized by its tendency to generate a new cluster, the value of its *α* parameter alters the predictive distribution. To establish an upper limit on the performance of the CRP, we used numerical optimization to determine *α* for each value of n. In addition, we also evaluated the performance of an agent with a fixed *α* = 1, which we believe is a more accurate reflection of a generalizer in an unknown environment. As expected, as we increase the structure of the sequences (lower entropy), the CRP advantage over a naïve guess increases [Fig S2, left, green line]. Similarly, the optimal value of *α* declines with the sequence structure, such that it is more advantageous to cluster as the sequence becomes more predictable [Fig S2, right]. However, this benefit is minimal for very unstructured sequences, and for fixed values of *α*, clustering for highly unstructured sequences (*H*(*X*) ≲ 1.45bits) yields worse CRP performance compared to a naïve guess.

**Supplementary Figure 2.**
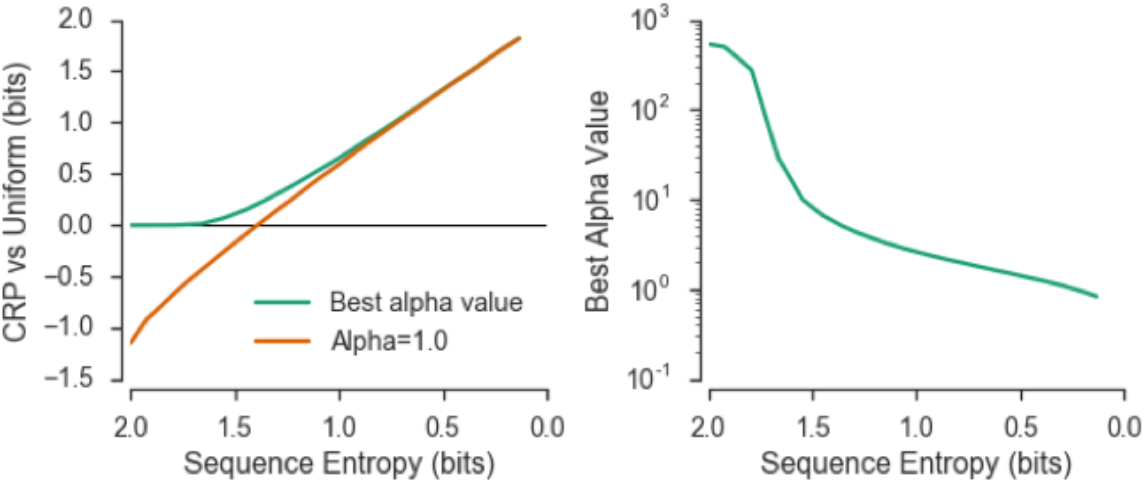
Performance of the CRP as a function of task domain structure. *Left*: Relative information gain of a naïve guess over the CRP as function of sequence entropy for a CRP with an optimized alpha parameter (green) or fixed at *α* = 1.0. *Right*: Optimized alpha value (log scale) as a function of sequence entropy.

## Footnotes

^1^Misclassification risk here is loosely defined: some misclassification errors are worse than others and misidentifying a cluster (and consequently, its MDP) may or may not result in a poor policy if the underlying MDPs are sufficiently similar. As such, the loss function in ecological settings can be a highly non-linear, domain specific function (as we demonstrate with the diabolical rooms problem).

